# Characterization and whole genome sequencing of *Saccharomyces cerevisiae* strains lacking several amino acid transporters: tools for studying amino acid transport

**DOI:** 10.1101/2024.12.03.626691

**Authors:** Unnati Sonawala, Aymeric Busidan, David Haak, Guillaume Pilot

## Abstract

*Saccharomyces cerevisiae* mutants have been used since the early 1980s as a tool to characterize genes from other organisms by functional complementation. This approach has been extremely successful in cloning and studying transporters, for instance, plant amino acid, sugar, urea, ammonium, peptide, sodium, and potassium were characterized using yeast mutants lacking these functions. Over the years, new strains lacking even more endogenous transporters have been developed, enabling the characterization of transport properties of heterologous proteins in a more precise way. Furthermore, these strains provide the added advantage of characterization of a transporter belonging to a family of proteins in isolation, and thus can be used to study the relative contribution of redundant transporters to the whole function. We focused on amino acid transport; starting with the yeast strain 22Δ8AA, developed to clone plant amino acid transporters in the early 2000s. We recently deleted two additional amino acid permeases, Gnp1 and Agp1, creating 22Δ10α. In the present work, five additional permeases (Bap3, Tat1, Tat2, Agp3, Bap2) were deleted from 22Δ10α genome in up to a combination of three at a time. Unexpectedly, the amino acid transport properties of the new strains were not very different from the parent, suggesting that these amino acid permeases play a minor role in amino acid uptake in our conditions. The inability to grow on a few amino acids as the sole nitrogen sources did not correlate with lower uptake activity, questioning the well-accepted relationship between lack of growth and loss of transport properties. Finally, in order to verify the mutations and the integrity of 22Δ10α genome, we performed whole-genome sequencing of 22Δ10α using long-read PacBio sequencing technology. We successfully assembled 22Δ10α’s genome *de novo*, identified all expected mutations and precisely characterized the nature of the deletions of the ten amino acid transporters. The sequencing data and genome will serve as a resource to researchers interested in using these strains as a tool for amino acid transport study.

## Introduction

Basic cell biology and metabolic pathways are well conserved among eukaryotic organisms, enabling using simpler eukaryotic organisms to study the function of proteins from more complex organisms. Heterologous expression of a gene allows to deduce the biochemical function and properties of the encoded protein and further understand its function [1]. The function of a gene can be better by expressing it in a yeast strain where the genes responsible for the same activity have been inactivated. Such a strain exhibits a phenotypic defect, which is reverted by the expression of the gene of interest, a method called functional complementation of the yeast mutant [2,3]. Many proteins from plants have been successfully studied by expression in baker’s yeast (*Saccharomyces cerevisiae*) [2,4–6].

Amino acids are critical for many cellular processes such as nitrogen homeostasis, protein synthesis, and nucleoside synthesis. Amino acid transporters, which mediate the translocation of amino acids across membranes, are bona fide components of metabolic pathways [7] and thus play a fundamental role in these functions. Several membrane proteins including amino acid transporters from both plants and animals have been characterized by expressing them in yeast cells [2]. Studying amino acid transporters by functional complementation of yeast transport mutants is achieved by testing the growth of yeast on a medium containing amino acids as the sole nitrogen source: the yeast can take up nitrogen (provided by the amino acid, [8]) only when the amino acid transport function is provided at the plasma membrane by the foreign gene. Alternatively, amino acid uptake of the expressed transporter can directly be screened and measured by determining the amount of radiolabeled amino acids taken up by cells when provided in the external medium [9,10].

*Saccharomyces cerevisiae* contains several endogenous amino acid permeases, 22 of which are localized to the plasma membrane [11]. They all belong to the APC (Amino acid-Polyamine- organo Cation) superfamily and are further divided into the YAT (Yeast Amino acid Transport), LAT (L-type Amino acid Transport), and ACT (Amino acid Choline Transporter) families [12]. Some of these transporters display broad specificity for amino acids, whereas others are more specific and transport only a few amino acids [8,13].

Functional complementation of yeast was first used to identify the amino acid transporters from the plant *Arabidopsis thaliana* in the early 1990s. The yeast strain JT16 lacking both the histidine permease (Hip1) and an enzyme required in the synthesis of histidine (His4) was used to screen for a plant amino acid transporter that could uptake histidine [14]. Around the same time, yeast strain 22574d (Fig 1) lacking a broad specificity permease (Gap1), a *γ*-aminobutyric acid permease (Uga4) and a high-affinity proline permease (Put4) was used to identify a plant amino acid transporter that was able to transport proline [15]. Both groups had simultaneously cloned the first secondary-active amino acid transporter, AAP1 (Amino Acid Permease 1) [16]. Several other plant amino acid transporters were identified in the second half of the 90’s by using complementation of these yeast mutants lacking their endogenous amino acid permeases [17]; see [18,10,19] for identification of other AAPs and related proline transporters, [20] for identification of cationic amino acid transporter (CAT1) and [21] for identification of lysine-histidine transporter (LHT1). The mutant yeast strain 22574d is still able to transport other amino acids apart from *γ*-aminobutyric acid (GABA) and proline by using the remaining endogenous amino acid permeases. To enable the study of lysine transport via heterologous expression, [22] further deleted three other yeast amino acid permeases responsible for transporting arginine (CAN1) and lysine (LYP1 and ALP1) along with an enzyme required for the biosynthesis of lysine (LYS2). This mutant yeast strain was named 22Δ6AAL (after the 6 deleted amino acid permeases in this strain and the biosynthesis enzyme Lys2). To study the transport of other amino acids, two other permeases were deleted in the 22Δ6AA background (HIP1 and DIP5 to disrupt histidine, and glutamate and aspartate transport respectively) leading to the strain 22Δ8AA [22] (Fig 1). 22Δ8AA is unable to grow on 5 amino acids as its sole nitrogen source supplied at 3 mmol.l^−1^, including the non-proteinogenic amino acids GABA and citrulline (Cit). This strain is still able to grow on 13 other amino acids. We previously reported deleting Agp1 and Gnp1 (shown to be necessary for the uptake of Threonine [23]) in the 22Δ8AA background to increase the number of amino acids whose transport can be studied, creating 22Δ10α [24]. 22Δ10α was found to be unable to grow on 10 additional amino acids as its sole nitrogen source compared to 22Δ8AA [24], probably because Agp1 is actually a broad specificity amino acid permease [25].

**Fig 1.**
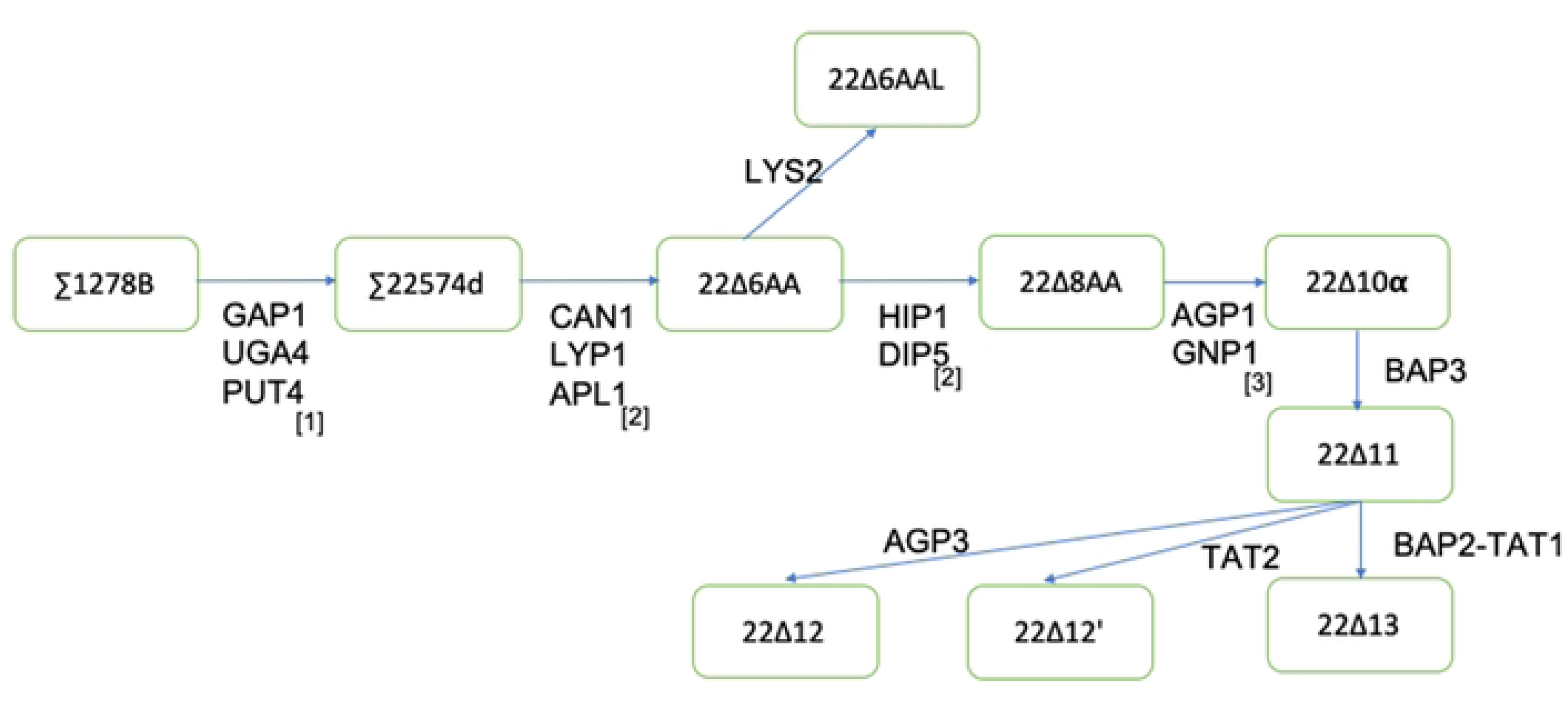
Schematic of genealogy leading to yeast strain 22Δ10α and its progenies. The major strain names are indicated in green boxes and the knock-out of genes encoding given amino acid permeases are noted beneath the arrows. 1: Jauniaux and Grenson, 1990 [38]. 2: Fischer et al. 2002 [22]. 3: Besnard et al. 2016 [24]

In the present work, we wanted to reduce the background transport for some aromatic amino acids by deleting five other endogenous amino acid permeases in the 22Δ10α background in three different combinations. We also tested if the resulting strains could be used to study transport at concentrations higher and lower than 3 mmol.l^−1^, which would be useful in characterizing low- or high-affinity plant amino acid permeases. Finally, the genome of 22Δ10α was sequenced and analyzed to identify any major chromosomal rearrangement that could have arisen during the successive gene deletion events.

## Material and Methods

### Yeast strains and manipulation

The present work proves that the mating type of 22Δ10α and hence its parent 22Δ8AA is MAT*a*; the genotype of both strains has thus been corrected in this Material and Method section. See Table 1 for the genotypes of the strains used in this work. Yeast cells were transformed using the lithium acetate method [26], and genes were deleted from the genome sequentially using the insertion of the kanMX cassette flanked by two loxP sites [27].

**Table 1.**
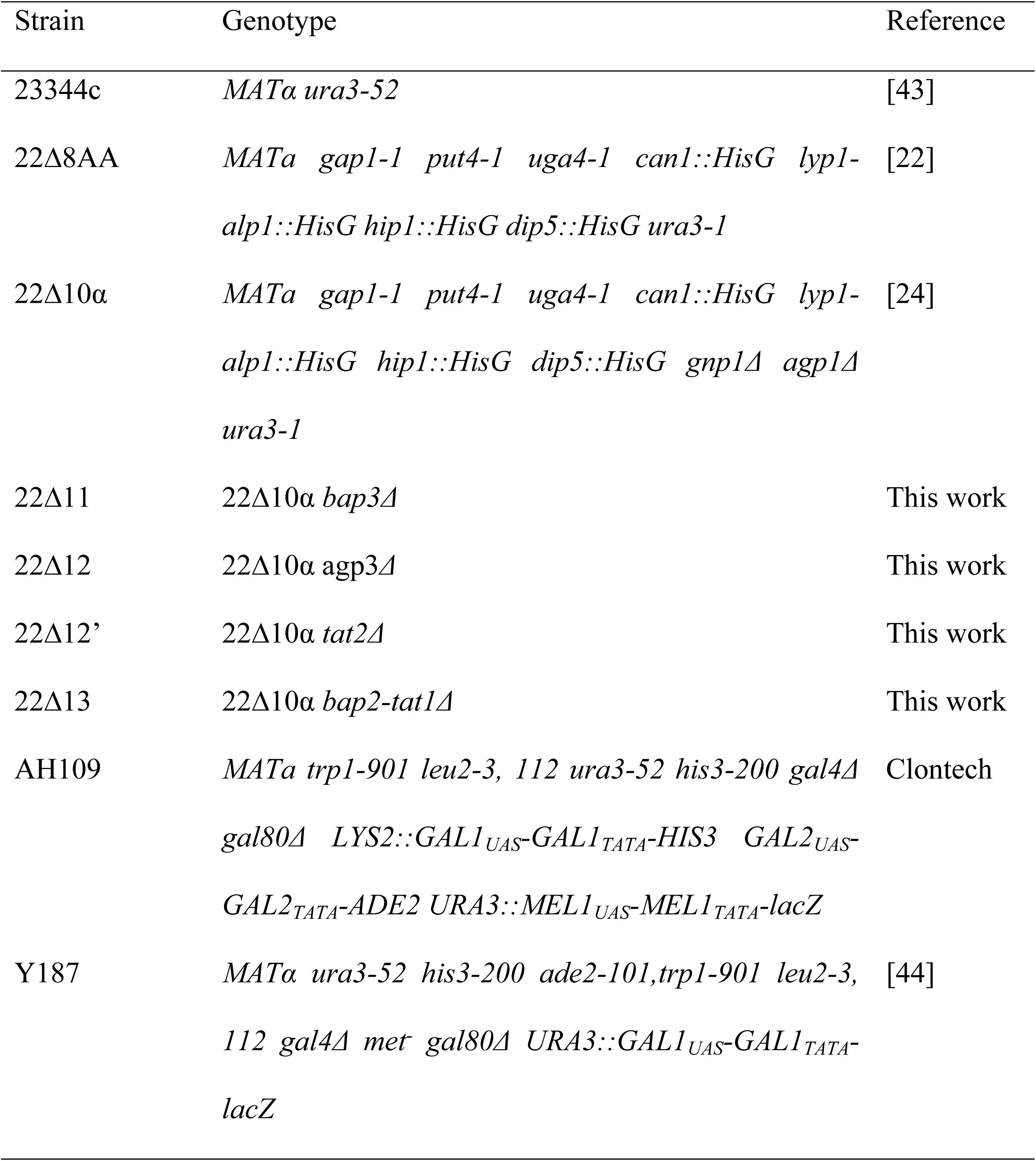
Yeast strains produced and used in this work.

### Cloning

ScGap1 was PCR-amplified from *Saccharomyces cerevisiae* genomic DNA, cloned into pDONRZeo using the Gateway technology (ThermoFisher Scientific), and moved to the destination vector pRS-Ura-Ws (a derivative of pRS416, in which the gateway-compatible expression cassette from pDR196-Ws was inserted).

### DNA extraction, sequencing and genome sequence analysis

Yeast cells were grown in 50 ml of YPDA medium (10 g/l yeast extract, 20 g/l bacto peptone, 20 g/l glucose, 80 mg/l adenine) (starting at OD of 0.1 at 600 nm) until they reached the exponential phase (two doublings), washed and resuspended in 3 ml of 1 M sorbitol. They were then converted to spheroplasts by adding 50 µl of a solution of 4 mg/ml Zymolyase 100T (United States Biological) in 50 mM Tris-HCl, pH 7.5, 1 mM EDTA and 50% glycerol, and incubating at 30^°^C with gentle shaking (70 rpm) for 1 hour, followed by a wash with 1 M Sorbitol. High molecular weight DNA was extracted from the pelleted spheroplasts using MagAttract HMW DNA kit (Qiagen) by following the manufacturer’s instructions. DNA was sequenced by Novogene on the PacBio platform. Continuous long reads were assembled *de novo* using canu (version 1.9) with the additional parameters of *minReadLength=5000* and *minOverlapLength=1000* [28]. Reference-guided contig rearrangement and scaffolding was performed using RagTag [29]. The genome sequence for the yeast strain was obtained from yeastgenome.org (version R-64-1-1) and used as a reference genome. Reads were mapped to S288C via minimap2 with the option *-ax map-pb*. NGMLR mapped reads and Sniffles (1.0.12) were used for calling structural variants [30]. Variant calls were restricted to reads with mapping quality >30 using the parameter *-q 30*. To get high confidence structural variants the output from Sniffles was filtered for variants with read support of greater than 30 reads and alternate allele frequency greater than 0.4. Alignments and variants were visualized with Integrative Genomics Viewer (version 2.7) [31].

### Yeast growth and radioactive uptake assays

Amino acid uptake assays were performed as described by [32] using ^3^H-labelled amino acids. Briefly, yeast cells were grown overnight in SD medium (6.7 g/l Yeast Nitrogen Base without amino acids (Difco), 20 g/l glucose, supplemented with amino acids but lacking uracil, pH 6.3) overnight at 30°C. They were then subcultured in 15 ml of SD medium at an OD of 0.1 and incubated at 30°C until they reached an OD of ∼0.5. Yeast cells were then pelleted at 2,500g and washed by resuspending in water and centrifuging at 2,500g. Washed cells were resuspended in uptake buffer (50 mM KH_2_PO_4_ and 600 mM sorbitol at pH 4.5) at an OD of 5. Fifty µL of these cells were aliquoted to be used for uptake per replicate of each amino acid and placed on ice until use. For the assay itself, 5 µL of 1 M glucose was added to 50 µL of the aliquoted cells in uptake buffer and incubated at 30°C in a thermal mixer for 5 minutes. Exactly 5 minutes later at 30°C, 55 µL of a mix of the unlabeled amino acid at the required concentration and 1 µCi of ^3^H-labelled amino acid in uptake buffer was added, and the resulting mix placed back on the thermal mixer for exactly 3 minutes. The cells were filtered using 24 mm Whatman filters (cat no. 1822-024) using a filtration manifold (DHI lab Filtration Manifold 10 x 20 ml, cat no. EQU-FM-10X20-SET): the cells were transferred to 5 ml of uptake buffer placed in the filtering device; the cells and buffer were drained through the filter under vacuum; 5 ml of uptake buffer was added and drained similarly. The filters were then added transferred to scintillation vials, filled with 4 ml of Ultima Gold XR (Revvity).

### Complementation assays

Yeast strains were grown overnight in SD medium and SC medium (1.7 g/l Yeast nitrogen base with ammonium sulfate, 20 g/l glucose pH 6.3), respectively. Yeast cells were diluted to the appropriate OD and 4 µL drops were laid on a minimum medium [33] supplemented with the specified concentration of the mentioned amino acids at 30°C.

### Yeast doubling time measurements

For measuring the growth rate and doubling of yeast strains, three independent 5 ml cultures in YPDA were grown overnight at 30°C, and used the next morning to start a subculture in either YPDA or SD (+Uracil) medium at OD of 0.1. The subculture was allowed to grow until an OD of about 0.5, and used to start 200 µL cultures in 96-well plates at an OD of about 0.05-0.1 in either YPDA or SD (+Uracil) as specified. Each biological replicate was also technically replicated twice on the 96-well plate. Synergy HTX plate reader (Biotek) was used for measuring the OD of the cultures with the following settings: temperature set at 30°C, measurements taken every 5 min at 600 nm, continuous orbital shaking at 559 nm (slow speed) for 15 hours. The growth curve from the data was fit to logistic equation and growth characteristics were measured using the Growthcurver package in R [34].

### Determination of mating type

Genomic DNA extracted from yeast grown overnight in YPDA was analyzed with specific oligonucleotides as described in [35]. For the mating test, 23344c, 22Δ10α, AH109 and Y187 were grown on solid YPDA and resuspended in 200 µl water. About 50 µl of each solution was mixed as described in the figure legend, and 10 µl were dropped on a YPDA plate. After overnight growth at 30°C, cells were resuspended in water, and streaked on SD medium with uracil as indicated in the figure legend and grown for two days at 30°C.

## Results and discussion

### Deletion of amino acid permeases in 22Δ10α

The work by Regenberg et al. [36] and Bianchi et al. [8] showed that the endogenous permeases Bap2, Bap3, Tat1 and Tat2 are important for the transport of aromatic amino acids (Phenylalanine, Tryptophan and Tyrosine), for transport of the branched-chain amino acids (Leucine, Isoleucine and Valine) and to a lesser degree alanine and glycine across the plasma membrane. Agp3 has been shown to transport leucine and becomes more important for yeast growth in low nutrient conditions or in yeast mutants lacking the broad-specificity permeases Gap1 and Agp1 [37]. Based on their importance in amino acid uptake by yeast cells, these five endogenous amino acid permeases were deleted from the genome of 22Δ10α in the following combinations: deletion of *BAP3* leading to 22Δ11, deletion of *BAP2* and *TAT1* in 22Δ11 background leading to 22Δ13, and deletion of *AGP3* or *TAT2* in the 22Δ11 background each leading to 22Δ12 or 22Δ12’ respectively (Fig 1). Several attempts were made to make further deletions in the 22Δ12 and 22Δ13 background but were unsuccessful.

### Characterization of the transport ability of the 22Δ11, 22Δ12, 22Δ12’ and 22Δ13 strains

To test if these deletions would reduce the background growth of the mutant strains on amino acids as sole nitrogen source, 23344C, 22Δ8AA, 22Δ10α, 22Δ12, 22Δ12’ and 22Δ13 cells were grown on minimum medium containing uracil with each amino acid added at 3 mmol.l^−1^ and mmol.l^−1^ as the sole nitrogen source. Compared to 23344c, 22Δ8AA was, as expected, unable to use Asp, Cit, GABA, Gly, Orn, Pro as a nitrogen source (Fig 2) [22]). Similarly, 22Δ10α was not able to use Ala, Asn, Gln, Gly, Ile, Leu, Met, Phe, Ser, Thr, Val Trp and Tyr as a nitrogen source in addition to Asp, Cit, GABA, Gly, Ornithine (Orn), Pro (Fig 2; S1 Fig) [24]). Compared to 22Δ10α, further gene deletions had little effect on the background growth of the cells, except for Arg when supplied at 3 mmol.l^−1^, and Asn, Gly and Met supplied at 12 mmol.l^−1^ (Fig 2 and S1 Fig). The ability to the strains to be functionally complemented by amino acid transporters expressed from a plasmid was tested by expressing the ScGap1 amino acid permease [38] in 22Δ10α, 22Δ11, 22Δ12’ and 22Δ13, comparing the growth to the cells transformed with an empty plasmid. For all tested amino acids and concentrations (0.5, 3, 9 or 12 mM), ScGap1 enabled a similar growth for each of the three strains, well above the background (S2 Fig). In this growth experiment, the background growth on Met was reduced further in 22Δ13 compared to the other three strains.

**Fig 2.**
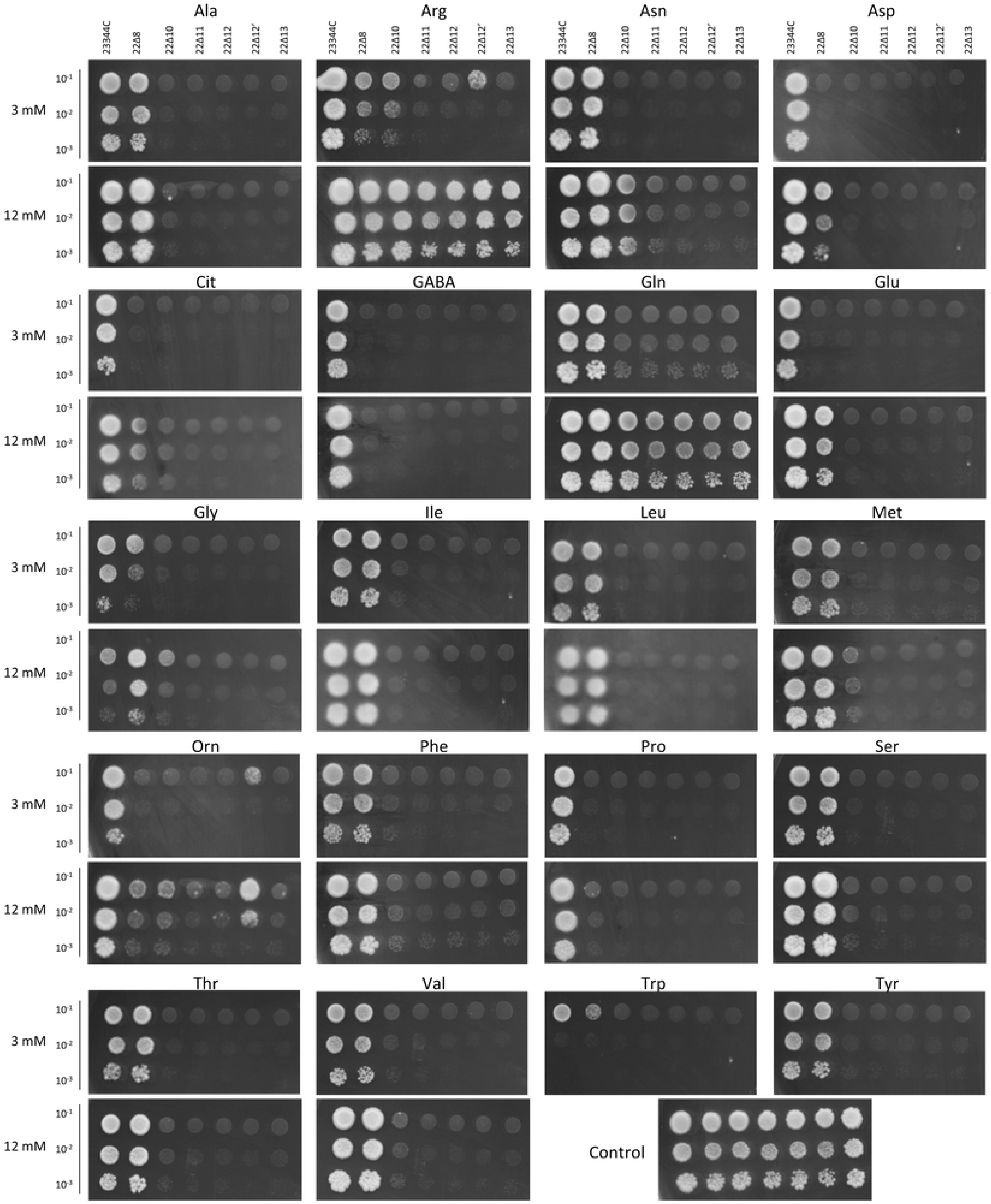
Growth comparison of 23344C, 22Δ10α, 22Δ11, 22Δ12, 22Δ12’ and 22Δ13 on amino acids as sole nitrogen source. OD for each strain was adjusted to 0.1, 0.01 and 0.001. Drops of 4 µl were aligned on minimum medium containing the labeled amino acid at 3 or 12 mM concentration as sole nitrogen source. The control medium contained 1.5 mM (NH4)2SO4. Growth was not tested on 12 mM Trp or Tyr because these amino acids are not stable at this concentration. Pictures were taken after 4 days at 30°C.

One of the limitations of the growth complementation assay above is its ineffectiveness when amino acids are supplied at very low concentrations (less than 0.5 mM) because the amount of amino acid/nitrogen transported is too low to support robust growth. A more suitable approach consists of directly measuring amino acid uptake by calculating the amount of radiolabeled amino acid taken up by the cells in a specific amount of time (typically 1-4 min). We thus tested whether the deletion of amino acid permeases in 22Δ10α lowered the background amino acid uptake when amino acids were supplied at low concentrations (3, 30 or 300 µM). For Asp, Gln, Met, Glu, Lys and Val supplied at any of the three concentrations, deletion of amino acid permease genes led to a reduction of uptake in 22Δ10α compared to the parental strain 23344c (Fig 3). Surprisingly, uptake of Leu, Trp, and Pro was not reduced in 22Δ10α compared to 23344c, despite a significant inability of this strain to grow on these three amino acids as sole nitrogen source (Fig 2). Overall, the background amino acid uptake was similar between 22Δ10α and 22Δ13, despite the three additional permease genes deleted (Fig 3).

**Fig 3.**
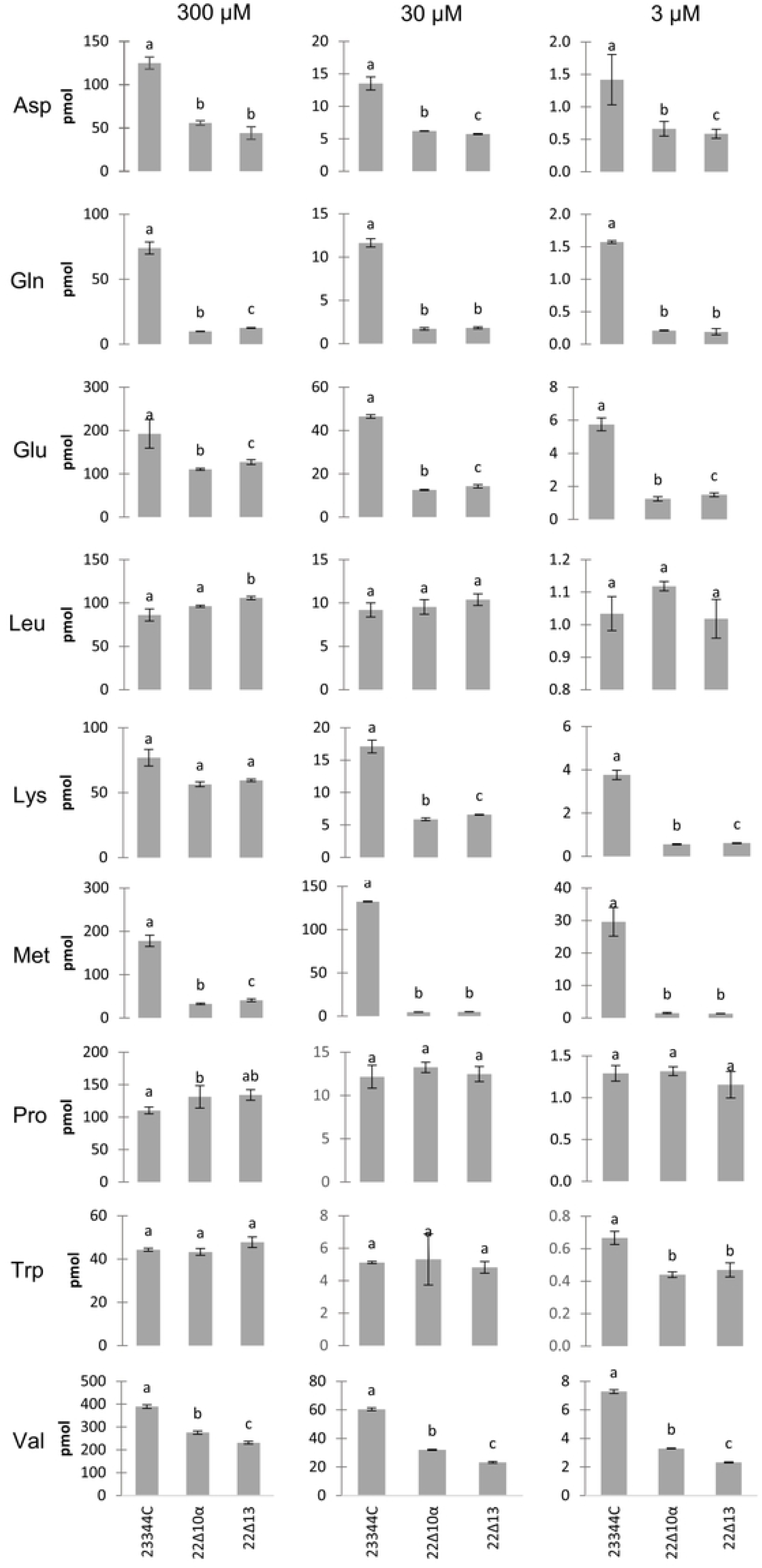
Uptake of radiolabeled amino acids by 23344C, 22Δ10α and 22Δ13 cells. Uptake of amino acids supplied at concentrations of 300 µM, 30 µM and 3 µM by parental yeast (23344c) and amino acid permease mutants (22Δ10α and 22Δ13) was measured after 3 minutes. Error bars represent standard deviation among technical replicates (n=3). Different letters indicate statistical significance at p<=0.05 according to t-test with Holm-Bonferroni correction for multiple comparisons.

Compared to the parental strain 23344c, 22Δ10α shows a deficiency in growth on and uptake of most of the proteogenic amino acids (this study and [24]). Surprisingly, further deletion of three extra endogenous amino acid permease genes (*BAP2*, *BAP3*, *TAT1*) creating 22Δ13 did not significantly improve these characteristics. The only improvement was observed when deleting BAP3 from 22Δ10α: 22Δ11 and its descendants showed deficiency in growth on 3 mM Arg as the sole nitrogen source when compared to 22Δ10α (Fig 2). Comparing growth and short-term uptake led to discrepancies: 22Δ10α and its descendants could not grow on Leu, Trp and Pro as the sole nitrogen source, but the uptake of these amino acids was not decreased in these strains compared to 23344c (except for Trp when supplied at 3 µM). It should be noted that such discrepancy may occur for other amino acids not evaluated in both assays (i.e. Ala, Arg, Asn, Cit, GABA, Gly, Ile, Lys, Orn, Phe, Ser, Thr, Tyr). The inability of 22Δ10α to grow on Pro as the sole nitrogen source is shared with its parent 22Δ8AA, supporting the validity of our results. The reason for this discrepancy is unknown but could result from several scenarios: To allow growth, the transporters need to be stable at the plasma membrane for hours/days after transfer to the complementation medium lacking ammonium; while direct uptake is measured right after growth in ammonium-rich medium. It is conceivable that the metabolic state of the yeast and the expression of the remaining set of amino acid transporters is different between the two conditions, possibly leading to this discrepancy. This phenomenon warrants further investigation because it casts doubt on the well-accepted assumption that observing the growth of a yeast strain by expression of a heterologous transporter means successful functional complementation. The slight reduction in background growth of 22Δ13 compared to the other strains suggests that Bap2 and Tat1 are mediating Met uptake when present at more than 3 mM outside of the cell.

### Sequencing and analysis of 22Δ10α genome

The mutant yeast strains starting from 22574d have been used to study plant amino acid transporters for several years. However, apart from the parent Σ1278b, none of these have been subjected to whole genome sequencing. Previous deletion studies, such as those for yeast deletion collection, have found that deleting yeast genes can cause off-target deletions or gene duplications, and sometimes chromosomal rearrangements [39,40]. Although rare, such collateral changes and off-target effects to the genome are possible. In order to aid future research with these mutant strains, we sequenced the 22Δ10α genome rather than progeny since they did not show much improvement in background growth. Using 8.12 Gbp of long-read PacBio sequencing data, 22Δ10α genome was assembled *de novo* using canu to an estimated depth of 600 [28]. When aligned to the reference genome (S288C), dotplot analysis showed coverage over all chromosomes further supporting the assembly. Furthermore, after scaffolding and polishing, no large duplications or translocations were identified (Fig 4a). Reference-guided separation of erroneously collapsed contigs and scaffolding was performed leading to a 12.87 Mbp assembly in 48 scaffolds, with 12.2 Mbp (95%) assembled in 16 scaffolds (Fig 4b, S3 Fig).

**Fig 4.**
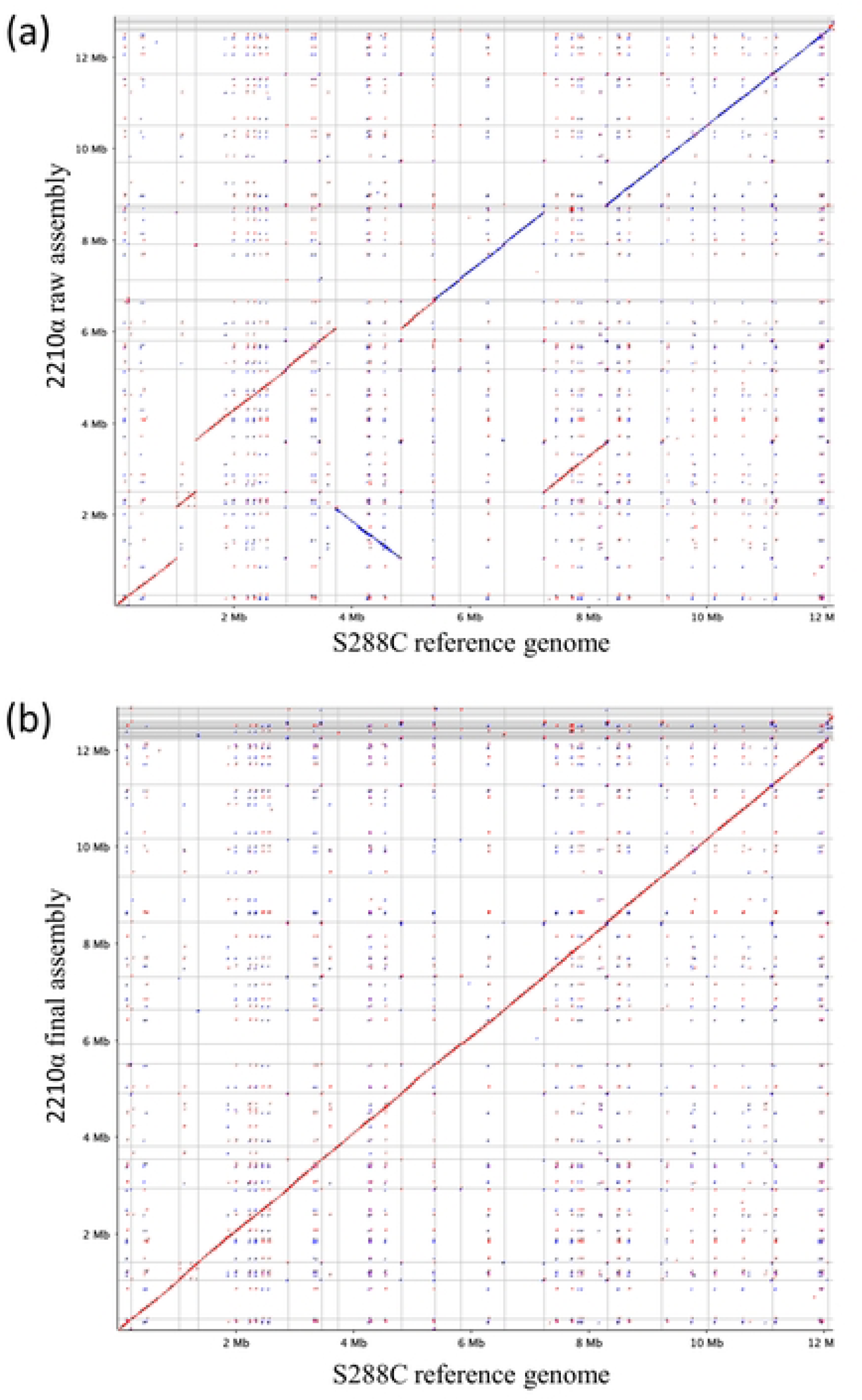
Whole genome alignment of 22Δ10α to S288C genome. Whole genome alignment of raw (a) and of the final (b) genome assembly of 22Δ10α against the S288C reference genome.

Structural variants (SV) were identified from 22Δ10α reads aligned to the reference genome (S288C) and used to identify structural variants (SV) using an SV caller specifically designed for long-read sequences with higher error rates. S288C genome was used for this purpose given its highly-contiguous chromosome-level assembly compared to the more fragmented Σ1278b assembly [41,42]. SV calls were filtered to get high-confidence variants (see methods), resulting in 221 nuclear SVs (S1 Data). The majority of the SVs were found to be related to transposable elements (TEs) and long-terminal repeats (LTRs) (data not shown). Since these structural variants were compared to the S288C reference genome many of these SVs could potentially stem from 22Δ10α parent strain Σ1278b and may not be specific to 22Δ10α.

We then identified the expected mutations of the 10 amino acid permeases genes, corresponding to loxP deletions, hisG insertions or other insertions (see below). In the case of insertions, reads spanning the expected gene were used to identify the insert using BLAST. All of the expected indels in the ten amino acid permease genes were correctly identified and confirmed (Table 2 and S4 Fig). The indel in *GAP1* resulted from an ∼4 kbp deletion replaced by a ∼6 kbp insertion of a Ty1 LTR retrotransposon; ∼6 kbp insertions of Ty1 elements were also found at the start of *PUT4* and within the *UGA4* loci [38]. *CAN1*, *HIP1, ALP1-LYP1* and *DIP5*, which were all deleted using the *hisG-URA3-neo-hisG* cassette [22], were found to have insertions ranging from ∼250 bp to 2.5 kbp left by the hisG cassette replacing the genes. *AGP1* and *GNP1*, which were deleted using the *loxP-kanMX-loxP* cassette [24], were both found to have 2 kb deletions abrogating the entire locus for each of these genes.

**Table 2.** Summary of the observed gene inactivations in 22Δ10α.

| Gene | Location of the InDel<sup>†</sup> | Deletion length (expected)<sup>‡</sup> | Insertion (inserted element)<sup>§</sup> |
| --- | --- | --- | --- |
| GAP1 | -1103bp | 4 kbp | 9 kbp (Transposon) |
| PUT4 | 5bp | - ¶ | 6 kbp (Transposon) |
| UGA4 | 1182bp | - ¶ | 6 kbp (Transposon) |
| CAN1 | 681bp | - ¶ | 250 bp (HisG scar) |
| ALP1-LYP1 | 835bp | 2.5 kbp | 1200 bp (HisG scar) |
| HIP1 | 1254bp | 250 bp | 1000 bp (HisG scar) |
| DIP5 | 538bp | - ¶ | 1200 bp (HisG scar) |
| AGP1 | -63bp | 2 kbp (2092 bp) | 34 bp (loxP) |
| GNP1 | -61bp | 2 kbp (2208 bp) | 34 bp (loxP) |

<sup>†</sup> Approximate bp from the ATG, with reference to ATG of ALP1 for ALP1-LYP1
<sup>‡</sup> Approximate size of the calculated deletion, using the S288C genome sequence as a reference.
<sup>§</sup> Approximate size of the DNA sequence replacing the gene deletion, with nature of the insertion. “Transposon”, transposon-related sequence; “HisG scar”, leftover sequence from the deletion approach [22]; “loxP” leftover loxP site from the deletion approach [24].
¶ No deletion was observed.

Genome sequencing also revealed that the mating locus of 22Δ10α was Mat*a*, while this strain has always been assigned with MAT*α* in the literature (see [22]). To determine 22Δ10α mating type, we genotyped the Mat locus by PCR and found that it was indeed Mat*a*, and we found that 22Δ10α mated with AH109 (MAT*a*) but not with Y187 (MAT*α*) (Fig 5). The mating type of 22Δ10α is therefore Mat*a* (note that the “α” of 22Δ10α is not related to its mating type, but its genealogy in the deletion procedure). The genealogy of 22Δ10α started from yeast strains published in 1970 and 1987 that were probably mated to create 22574d [38], the direct progenitor of 22Δ6AA, 22Δ8AA [22] and finally 22Δ10α [24]. It is thus unknown when this mistake occurred and if any of the parental strains were tested for mating type at any time.

**Fig 5.**
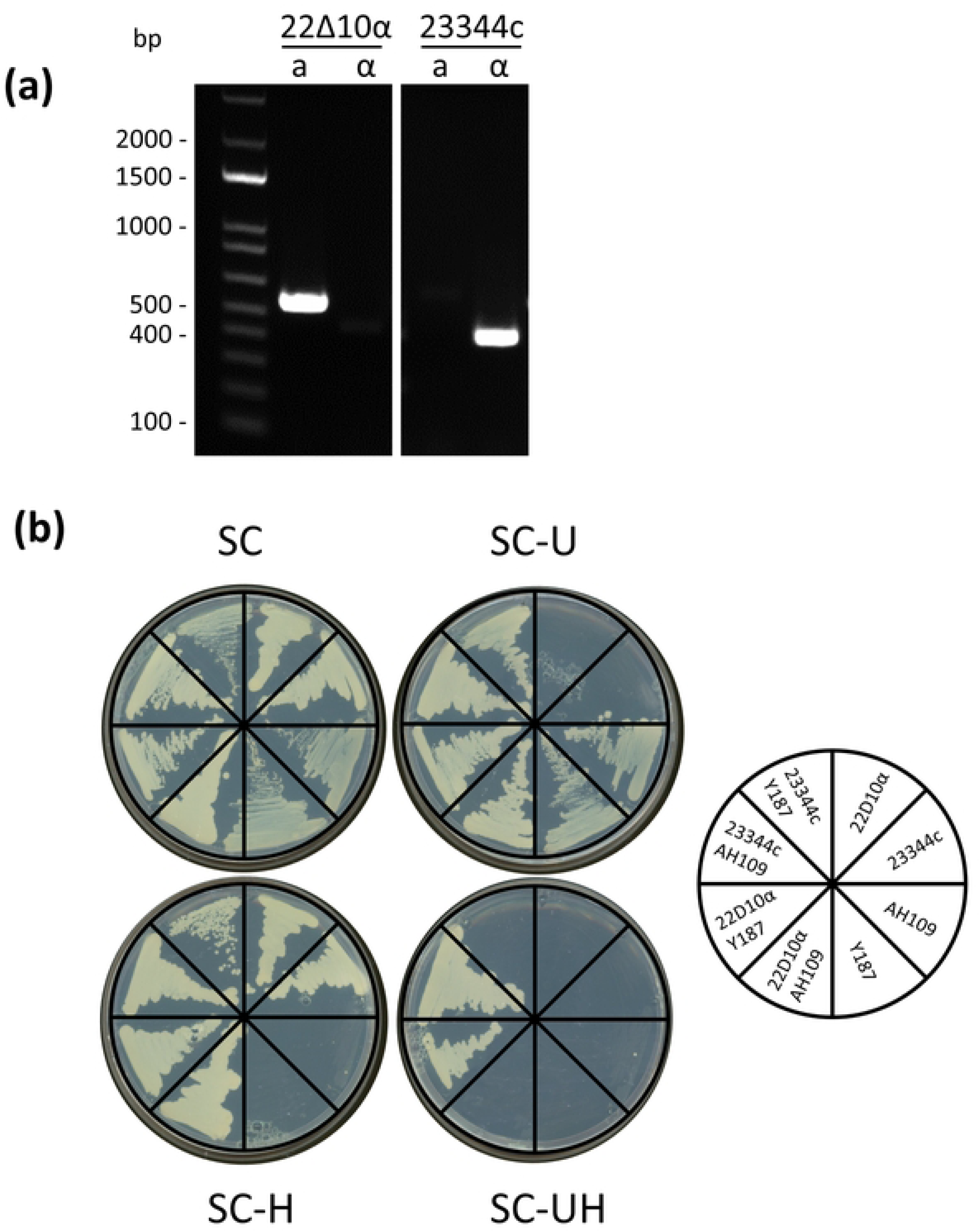
Mating type genotype and assay of 22Δ10α. (a) Genotyping PCR of 22Δ10α and 23344c strains with oligo pairs that correspond to the *Mat*a (a) or *Mat*α (α) mating type. Expected sizes are *Mat*a: 544 bp and *Mat*α: 404 bp. (b) Mating assay between 23344c and 22Δ10α with Y187 and AH109. Cells were grown overnight in liquid YPDA, mixed and dropped on YDPA plate overnight for mating and streaked on SC medium lacking Uracil (-U), His (-U) and both (-UH). Diploid cells, resulting from mating of cells of different mating types should be both Uracil and His prototroph.

### Analysis of growth speed of 22Δ10α, 22Δ11 and 22Δ13

While working with 22Δ11 and 22Δ13, these two strains were observed to grow appreciably slower compared to 22Δ10α and sometimes led to poor transformation efficiency. The growth rate of these strains and the corresponding doubling time was calculated for their growth in nutrient-rich YPDA medium and the minimal SD medium (S5 Fig). In the nutrient-rich YPDA medium 22Δ11, 22Δ12, 22Δ12’ and 22Δ13 took 25-30 minutes more to double compared to the wild-type (23344C) yeast strain. The impeded growth was magnified when these mutants were grown in minimal SD medium with 22Δ11, 22Δ12 and 22Δ13 doubling up to 40-50 minutes slower than 23344C. Later analysis of sibling colonies kept frozen during the generation of 22Δ11 showed that the slow growth happened not during but in the step following the cre-lox recombination used to remove the KanMx cassette. This step corresponded to curing of the pSH47 plasmid, and about half of the tested colonies displayed the same long doubling time as 22Δ11, while the other grew similar to 22Δ10α. Genotyping of the deletion sites of *GNP1*, *AGP1* and *BAP3* did not provide evidence of any genome rearrangement in any of the 22Δ11 sibling or descendants. This data suggests that the longer doubling time is not caused by deletion of *BAP3*. Since 22Δ12, 22Δ12’ and 22Δ13 are descendants of 22Δ11, they all grow slower than 22Δ10α, but this does not affect their potency as a tool for characterizing amino acid transport and transporters.

## Data Availability

The sequencing data and final genome assembly have been submitted under BioProject ID PRJNA862461

## Acknowledgements

The authors thank Nima Trivedi for help in gene deletion and initial testing of the yeast strains on minimum media. This work was supported by the National Science Foundation of USA (Grant IOS-1353366 to GP) and the Hatch Program of the National Institute of Food and Agriculture of USA (VA-135908 for GP) and the Virginia Agricultural Experiment Station.

## Supporting information

**S1 Figure. Growth assay comparing growth of 23344C, 22Δ10α, 22Δ11, 22Δ12, 22Δ12’ and 22Δ13 cells on given amino acid as sole nitrogen source.** Yeast cells were grown overnight in synthetic defined (SD) medium supplemented with uracil. OD for each strain was adjusted to 0.1, 0.01 and 0.001. Drops of 4µL were aligned on minimum medium containing labeled amino acid at 3 (**A**) or 12 mmol/l (**B**) as sole nitrogen source. Pictures were taken after 2.5 days growth at 30°C.

**S2 Fig. Functional complementation assay of 22Δ10α, 22Δ11, 22Δ12’, and 22Δ13 yeast strains.** 22Δ10α, 22Δ11, 22Δ12’, and 22Δ13 yeast strains were transformed with empty vector pRS-Ws or pRS-Ws vector containing the cDNA of GAP1 (General Amino acid Permease I, YKR039W). Yeast cells were grown overnight in selective medium. OD for each culture was adjusted to 1, 0.1 and 0.01. Drops of 5 µL were aligned on minimum medium containing amino acids at 0.5, 3, 9 or 12 mmol/l as sole nitrogen source, and grown at 30°C.

**S3 Fig. Integrated Genome Viewer images of indels in 22Δ10α** (a-i)22Δ10α PacBio sequencing reads were aligned to the S288C reference genome and were loaded in IGV (Robinson et al. (2011)) along with the reference genome and its annotation. In mapped reads, deletions are indicated by a black line and insertions by purple regions (numbers indicated length of deletion or insertion). When a read is clipped by more than 100 bp the end of that read is marked in red. Indels of less than 30bp are not labeled.

**S4 Fig. Assembly statistics of 22Δ10α genome.** 22Δ10α assembly statistics (left); plot cumulative length of the assembly vs. number of scaffolds (right)

**S5 Fig. Doubling time of yeast strains in YPDA and SD medium.** Yeast strains were grown in YPDA (a) or SD supplemented with URA (b) medium, and their ODs measured using a plate reader, every 5 min for 15 hours. The data were fitted to a standard form of logistic equations to get the growth characteristics including doubling time. Each boxplot the distribution of represents six data points per strain. * p-value<0.05

**S1 Data. Vcf file of the sequence variants obtained from genome comparison of 22Δ10α and S288C.**

